# It takes two: Aberrant repair and low-grade inflammation characterizes bronchiolitis obliterans syndrome after lung transplantation in serum proteomic analysis

**DOI:** 10.1101/2025.01.13.632807

**Authors:** Eline A. van der Ploeg, Alen Faiz, Greta J. Teitsma, Alejandro Sánchez Brotons, Natalia Govorukhina, Jannie M.B. Sand, Diana J. Leeming, Barbro N. Melgert, Peter Horvatovich, Janette K. Burgess, C. Tji Gan

## Abstract

**Aim:** The obstructive phenotype of chronic lung allograft dysfunction, bronchiolitis obliterans syndrome (BOS), is diagnosed after lung transplantation (LTx) when irreversible airway obstruction is already present. This study aimed to investigate the fibrotic response and inflammation signals in serum of BOS patients.

**Methods:** LTx patients transplanted at the University Medical Center Groningen between 2004 and 2017 were screened. Nineteen patients with BOS were selected and matched to 19 non-BOS patients. Only patients for whom lung function and longitudinal serum samples post-LTx were available were included. Enzyme-linked immunosorbent assays were performed for neoepitopes of collagen types I, III, and VI and osteoprotegerin (OPG) in serum. Additionally, serum samples were analyzed by label free liquid chromatography with tandem mass spectrometry proteomics analysis.

**Results:** Collagen neoepitopes did not differ significantly between BOS and non-BOS patients at any timepoint. OPG was significantly higher in non-BOS compared to BOS six months before BOS onset (p<0.04). In proteomics analysis, proteins indicating cell repair and proliferation, namely human type II keratin-6 and centromere protein F (both FDR<0.1), were significantly lower three months before BOS onset in BOS compared to non-BOS patients. C-reactive protein (FDR<0.05) and SERPINA3 (FDR<0.05) amongst others, were higher in end-stage BOS compared to non-BOS patients.

**Conclusion:** Differences in expression of proteins that reflect the complex interplay between fibrosis and inflammation in BOS were identified. These proteins should be investigated and validated in larger cohorts and may aid in expanding knowledge about the development of BOS.

## • Introduction

Lung transplantation (LTx) is a life-saving treatment for several lung diseases like chronic obstructive pulmonary disease and idiopathic pulmonary fibrosis^1^. Annually, 3,800 LTx are performed worldwide^2^, however, long-term survival is restricted mainly by development of chronic lung allograft dysfunction (CLAD). CLAD affects half of all patients five years after LTx, causing major morbidity and mortality^3,4^. Two main forms of CLAD can be identified: restrictive allograft syndrome (RAS) and bronchiolitis obliterans syndrome (BOS)^5^. RAS is characterized by restrictive pulmonary function reflected by rapid decline of forced expiratory volume in one second (FEV1) and total lung capacity (TLC) as well as persistent pulmonary opacities on computed tomography. In BOS, the most common form of CLAD, obstructive pulmonary function is the hallmark for diagnosis in absence of other factors such as infection. Patients develop a persistent decline in FEV1 and FEV1/forced vital capacity (FVC) ratio representing irreversible damage^5^. BOS is pathologically characterized by obliterative bronchiolitis that causes progressive obliteration of small airway lumina in the donor lung^5^. The patchy occurrence of the fibrotic lesions complicates early pathological diagnosis^3^. The pathogenesis of BOS remains only partially understood. Allo-immune dependent as well as independent factors, such as infection and gastro-esophageal reflux, are known to contribute to the complex pathogenesis^6,7^. Repeated damage to the airway induces recruitment of neutrophils as well as (myo)fibroblasts, amongst others. Epithelial to mesenchymal transition, tissue remodeling and excess production of extracellular matrix, including accumulation of collagens, collectively progress fibrotic obliteration of the airway^8,9^. Changes in extracellular matrix composition can be seen in biopsies from LTx patients before BOS onset compared to controls, highlighting the importance of the extracellular matrix in BOS^10^. Earlier research in idiopathic pulmonary fibrosis (IPF) showed increased levels of N-terminal propeptide of type III collagen (PRO-C3), N-terminal propeptide of type VI collagen (PRO-C6), a fragment of type I collagen released by MMP (C1M) and a fragment of type VIα1 collagen released by MMP-2 (C6M) in serum, all markers for collagen types I, III and VI turnover, that related to progression of disease^11–15^. Also, osteoprotegerin (OPG), known for its role in matrix turnover and instigating fibrogenesis through upregulation of TGF-β1^16^, was described to be increased in human and mouse fibrotic lung tissue and to associate with progressive fibrotic disease^17^. Since excess production of extracellular matrix, particularly accumulation of collagens, is a hallmark of BOS, we hypothesized that these blood biomarkers could also reflect disease in BOS, possibly before onset of lung function decline. We also hypothesized that hypothesis-free protein profiling through mass spectrometry of serum of LTx BOS and non-BOS patients could provide novel insights into the underlying fibrotic molecular and cellular processes driving BOS. Therefore, we performed label free proteomics on serum of BOS and matched non-BOS patients including samples before and after BOS onset. To our knowledge, label free serum proteomics have not been performed in BOS before.

## 2.1 Methods

### 2.1.1 Study design and patient selection

LTx patients of eighteen years and older who underwent bilateral transplantation between 2004 and 2017 in the University Medical Center Groningen, the Netherlands, were screened for eligibility. We started screening from 2004 onwards because of the introduction of tacrolimus. Nineteen patients who developed BOS and who all progressed to BOS stage three ascertained on International Society of Heart and Lung Transplantation Guidelines^18^ were selected based on availability of at least three longitudinal serum samples (Figure 1). Patients with mixed CLAD phenotypes were excluded. BOS patients were matched with 19 non-BOS LTx patients. Patients were matched for sex, age at LTx, disease necessitating LTx, type of immunosuppression used and total storage time of samples. At the timepoints of data and serum collection other causes of lung function decline were excluded. All patients received standardized immunosuppression, pulmonary function follow up and clinical management according to the local LTx protocol. Patients provided written informed consent for use of material. The study was approved by the medical ethics committee of the University Medical Centre Groningen (METc 2021/610, research register number: 202000737).

**Figure 1.**
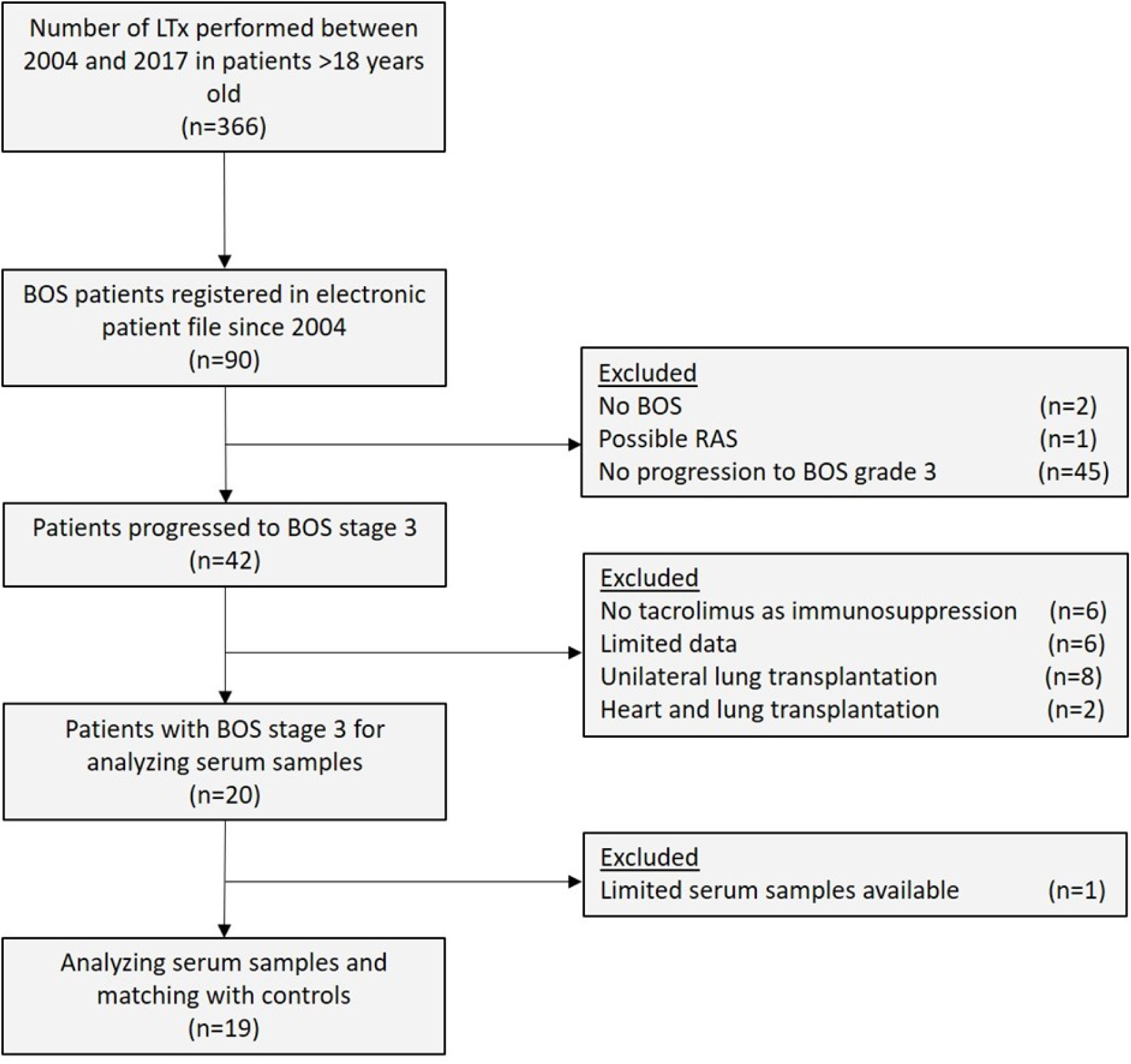
Flowchart indicating patient selection for BOS patients matched to non-BOS patients. Abbreviations: LTx: lung transplantation; BOS: bronchiolitis obliterans syndrome; RAS: restrictive allograft syndrome; n: number.

### 2.1.2 Sample collection

At the timepoints of data collection, other causes of lung function decline were excluded. Timepoints for serum sample collection of patients were 12, six, and three months before BOS onset (BOS stage 1), at BOS stage 1, at BOS stage 2 and BOS stage 3. BOS grading was defined according to the ISHLT guideline^19^. For non-BOS patients, dates of the serum sample collections were matched in relation to time since LTx date compared to the BOS patients.

Serum samples from 19 BOS and 19 non-BOS patients were analyzed by multiple ELISAs for collagen neoepitope testing for all time points available. Serum samples of 18 BOS and 16 non-BOS patients were analyzed by label free proteomics analysis. (Figure 2). Proteomics was performed on samples at timepoints minus three months before BOS onset, BOS stage 1 and BOS stage 3 samples in BOS and non-BOS patients. Missing samples were not replaced (Supplementary Table 1 and 2).

**Figure 2.**
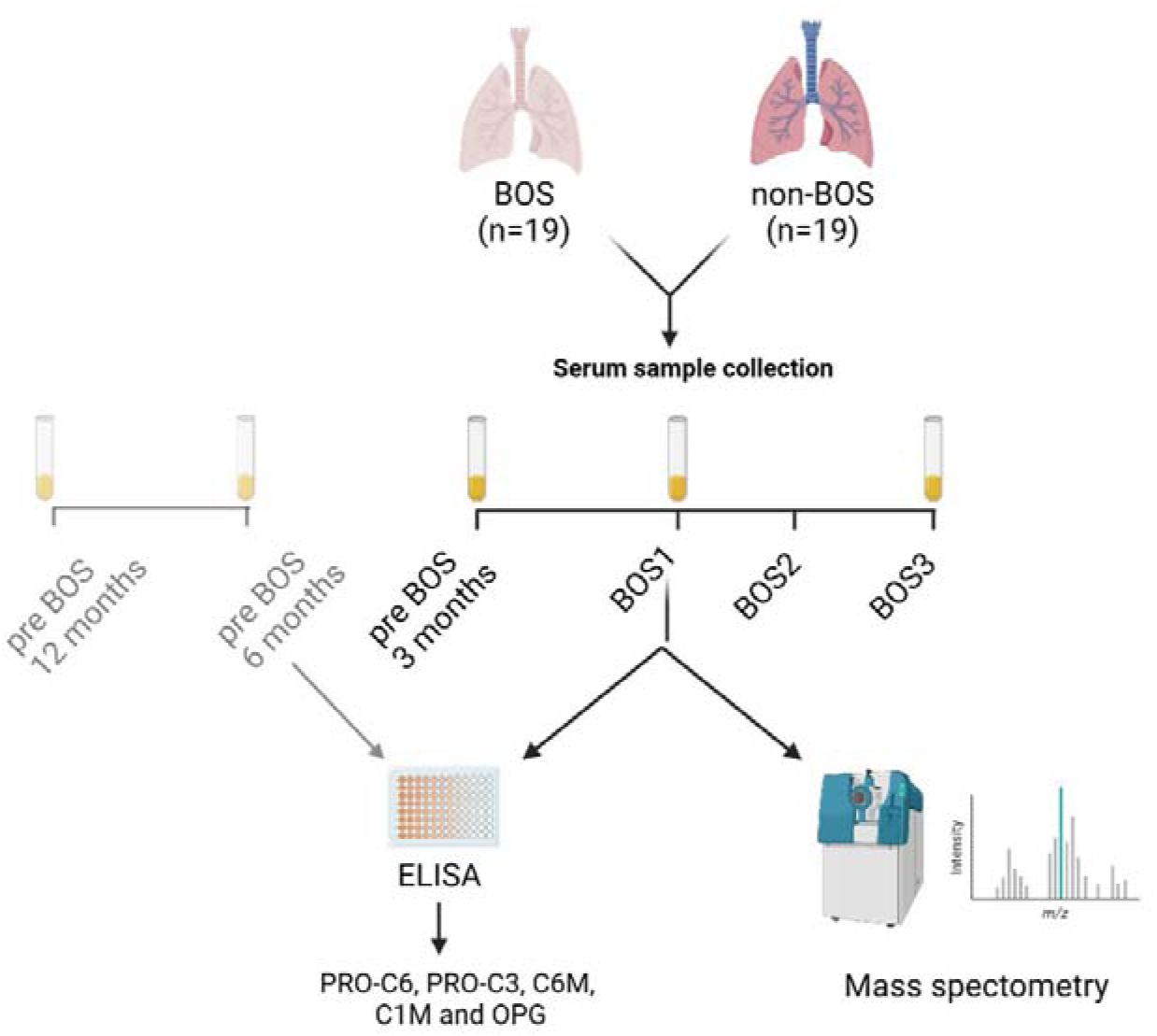
Flow diagram of methods used for analysis of serum samples of BOS and non-BOS patients. For OPG and collagen fragment analysis, samples were collected at 12, six, and three months before onset of BOS, at BOS stages 1 and 3 (BOS n=19 and non-BOS n=19). Samples from BOS and non-BOS patients were collected at three months before BOS onset, BOS stage 1 and 3 for analyses with mass spectrometry (BOS n=18, non-BOS n=16). Abbreviations: BOS: bronchiolitis obliterans syndrome; BOS1: BOS stage 1; BOS2: BOS stage 2; BOS3: BOS stage 3; PRO-C3: type III collagen formation; PRO-C6: type VI collagen formation; C1M: type I collagen degradation; C6M: type VI collagen degradation; OPG: osteoprotegerin. Figure composed with Biorender.

### 2.1.3 OPG and collagen neoepitopes

OPG was measured in serum using a human OPG DuoSet ELISA kit (R&D Systems, Minneapolis, US) according to the instructions provided by the manufacturer. Collagen neoepitopes C1M, C6M, PRO-C3, and PRO-C6 were measured in serum using neoepitope specific competitive ELISAs developed and validated by Nordic Bioscience (Herlev, Denmark) according to previously published protocols: C1M and C6M quantify type I and VI collagen degradation fragments, respectively, generated by matrix metalloproteinases and were measured using the nordicC1M^TM^ and nordicC6M^TM^ assays^20,21^. PRO-C3 and PRO-C6 quantify type III and VI collagen formation, respectively, and were measured using the nordicPRO-C3^TM^ and nordicPRO-C6^TM^ assays^22,23^.

### 2.1.4 Proteomics

The proteomic analysis was performed on serum samples after depletion of the most abundant proteins such as albumin. Proteins were digested with the use of trypsin. Using liquid chromatography with tandem mass spectrometry (LC-MS/MS) analysis, samples were analyzed, and data retrieved. Peptide and protein identification was performed using a false discovery rate of 1% at peptide-spectrum matching, peptide and protein levels. Human proteome from Swissprot was used for identification of proteome, after which quantitative processing was performed with PASTAQ^24^ (See Figure 3. For a detailed description of proteomics methodology see Supplement 1).

**Figure 3.**
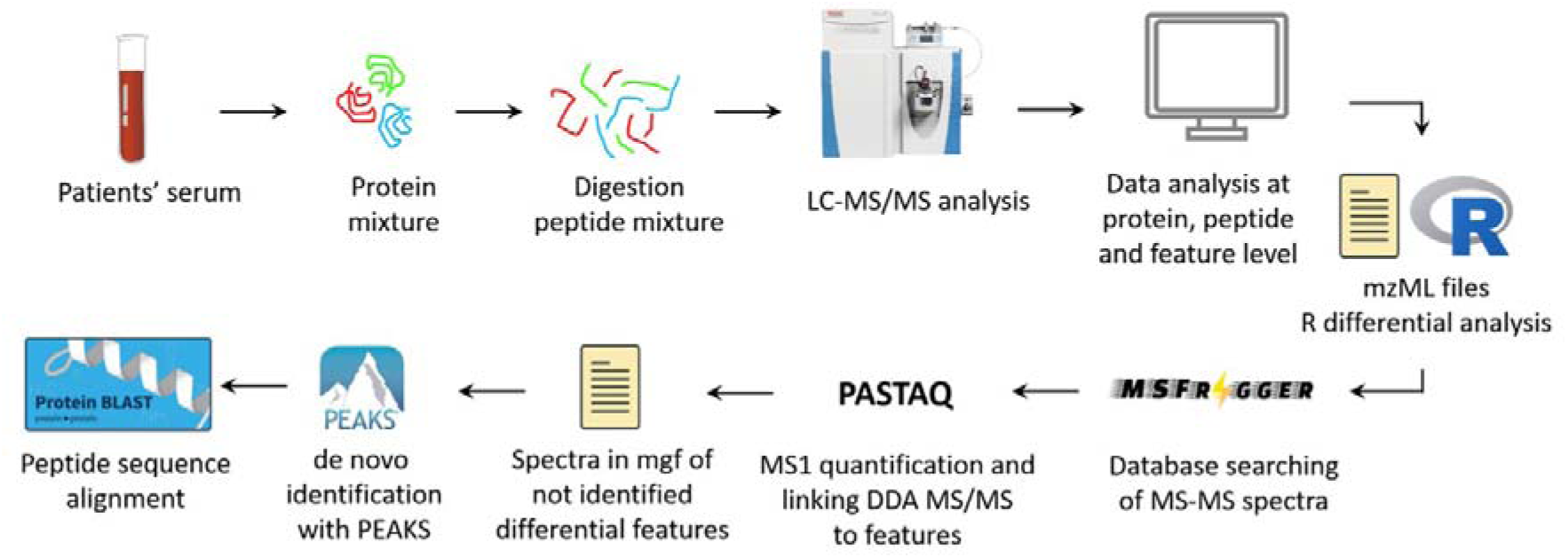
Flow diagram for proteomics analysis on serum samples of BOS and non-BOS patients. Patients’ serum were depleted and analyzed by LC-MS/MS analysis. Data retrieved were processed with R, MSfragger and PASTAQ, after which *de novo* identification for proteins with PEAKS was performed. Abbreviations: LC-MS/MS: liquid chromatography with tandem mass spectrometry; MS-MS: tandem mass spectrometry; DDA: data dependent analysis; mgf: mascot generic format.

### 2.1.5 C-reactive protein collection

C-reactive protein (CRP) is routinely measured using turbidimetry during follow up visits in the University Medical Center Groningen. For validation of proteomics results, CRP data was retrospectively collected from electronic patients’ files.

### 2.1.6 Statistical Analyses

Statistical analyses were performed using RStudio 2021.09.1 Build 372 and IBM SPSS version 28.0. Differential expression analyses were performed using the Limma R package across peptide, protein and feature data sets. All proteomic analyses were corrected for sex, age, and alpha-1 antitrypsin deficiency (AATD), to check for associations between these variables, BOS, and protein levels. False discovery rate for peptides was set at 0.05. For protein and feature level the false discovery rate was set at 0.1 to allow for identification of new proteins. Data for proteomics analysis were log transformed. Continuous data is represented as mean ± standard deviation (SD) or median [interquartile range] depending on normality. Differences in patient characteristics between BOS and non-BOS patients were analyzed by Mann-Whitney U test for continuous not normally distributed data and Chi-square for categorical data. Independent samples T-test was used to analyze continuous normally distributed data. ELISA results between groups were analyzed at each time point with Mann-Whitney U test for matched data and Wilcoxon signed rank test for paired data within groups. A p-value < 0.05 was considered significant.

## 3.1 Results

### 3.1.1 Patient characteristics

The median age at LTx was 55 years, with chronic obstructive pulmonary disease being the most common reason for LTx, followed by alpha-1 anti-trypsin deficiency (Table 1). Patients developed BOS 2.8 [1.9-5.8] years after LTx. There was no significant difference in time between LTx and serum samples collected between BOS and non-BOS patients for all time points (p>0.2, supplementary table 3). FEV1 and FVC in liters, as well as in percentage predicted, differed significantly from onset of BOS diagnosis in BOS compared to non-BOS patients, supporting diagnosis (Supplementary figure 1). FEV1 as percentage from baseline post LTx did not differ between BOS and non-BOS patients at timepoints before BOS onset, however a steady decline was observed from BOS onset in BOS group compared to non-BOS patients reflecting BOS diagnosis (Figure 4).

**Table 1.**
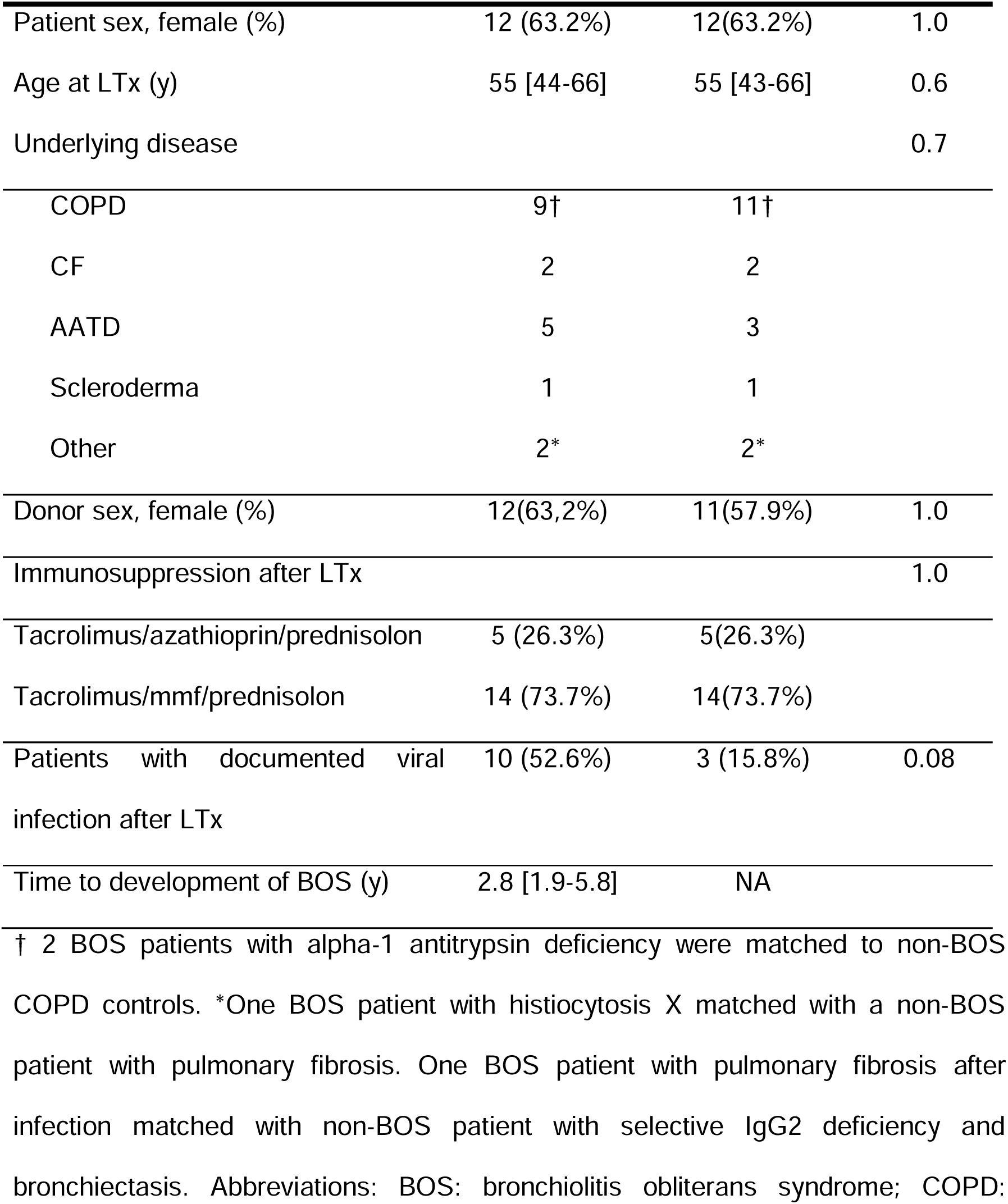
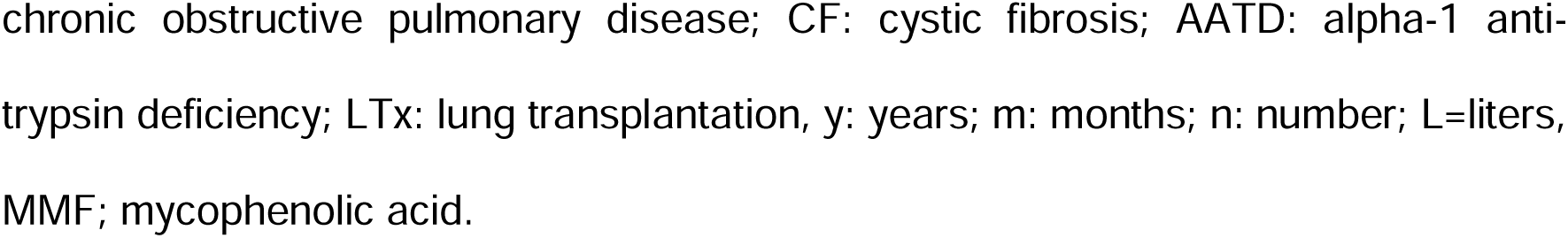
Patient characteristics of BOS and non-BOS patients.

|  | **BOS (n=19)** | **Non-BOS (n=19)** | **p-value** |
| --- | --- | --- | --- |
| Patient sex, female (%) | 12 (63.2%) | 12(63.2%) | 1.0 |
| Age at LTx (y) | 55 [44-66] | 55 [43-66] | 0.6 |
| Underlying disease |  |  | 0.7 |
| COPD | 9† | 11† |  |
| CF | 2 | 2 |  |
| AATD | 5 | 3 |  |
| Scleroderma | 1 | 1 |  |
| Other | 2* | 2* |  |
| Donor sex, female (%) | 12(63,2%) | 11(57.9%) | 1.0 |
| Immunosuppression after LTx |  |  | 1.0 |
| Tacrolimus/azathioprin/prednisolon | 5 (26.3%) | 5(26.3%) |  |
| Tacrolimus/mmf/prednisolon | 14 (73.7%) | 14(73.7%) |  |
| Patients with documented viral infection after LTx | 10 (52.6%) | 3 (15.8%) | 0.08 |
| Time to development of BOS (y) | 2.8 [1.9-5.8] | NA |  |

† 2 BOS patients with alpha-1 antitrypsin deficiency were matched to non-BOS COPD controls. \*One BOS patient with histiocytosis X matched with a non-BOS patient with pulmonary fibrosis. One BOS patient with pulmonary fibrosis after infection matched with non-BOS patient with selective IgG2 deficiency and bronchiectasis. Abbreviations: BOS: bronchiolitis obliterans syndrome; COPD:
chronic obstructive pulmonary disease; CF: cystic fibrosis; AATD: alpha-1 anti-trypsin deficiency; LTx: lung transplantation, y: years; m: months; n: number; L=liters, MMF; mycophenolic acid.

**Figure 4.**
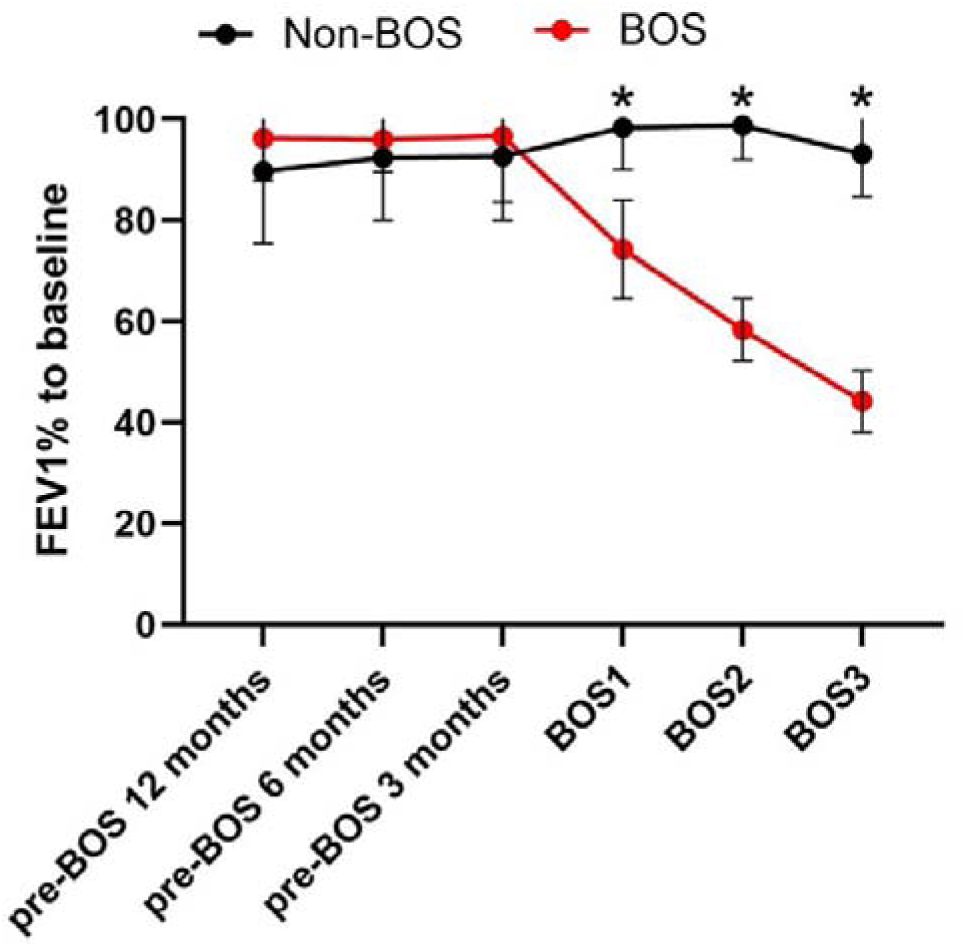
Forced Expiratory Volume in one second in percentage compared to baseline measurements for BOS and non-BOS patients. Baseline measurements were defined as the average of the best two measurements post-lung transplantation with at least three weeks separation. At BOS stage 1, 2 and 3, FEV1 percentage compared to baseline was significantly different between BOS and non-BOS patients when analyzed with unpaired T-test (BOS stage 1: 74.3±9.8 vs 95.5±12.7, p<0.001, BOS stage 2: 58.4±6.2 vs 98.8±6.8, p<0.001, BOS stage 3: 44.2±5.9 vs 93.2±8.4, p<0.001). All error bars reflect mean ± standard deviation. Abbreviations: BOS; bronchiolitis obliterans syndrome: BOS1; BOS stage 1: BOS2; BOS stage 2: BOS3; BOS stage 3. * p-value <0.05.

### 3.1.2 Fibrosis markers osteoprotegerin and collagen neoepitopes are not associated with BOS

OPG serum levels were significantly lower in patients developing BOS 6 months before BOS onset compared to non-BOS patients (1932 pg/ml vs 3685 pg/ml, p=0.04). However, OPG levels were not different at other timepoints (Figure 5). For the collagen neoepitopes, neither C1M, C6M, PRO-C6 or PRO-C3 serum levels differed in BOS patients compared to non-BOS patients (p>0.05 across all time points; Figure 6). However, a trend towards higher C1M levels in BOS patients compared to non-BOS patients was found for all timepoints (Figure 6A). ROC analysis did not show predictive value for any of the timepoints for C1M (data not shown).

**Figure 5.**
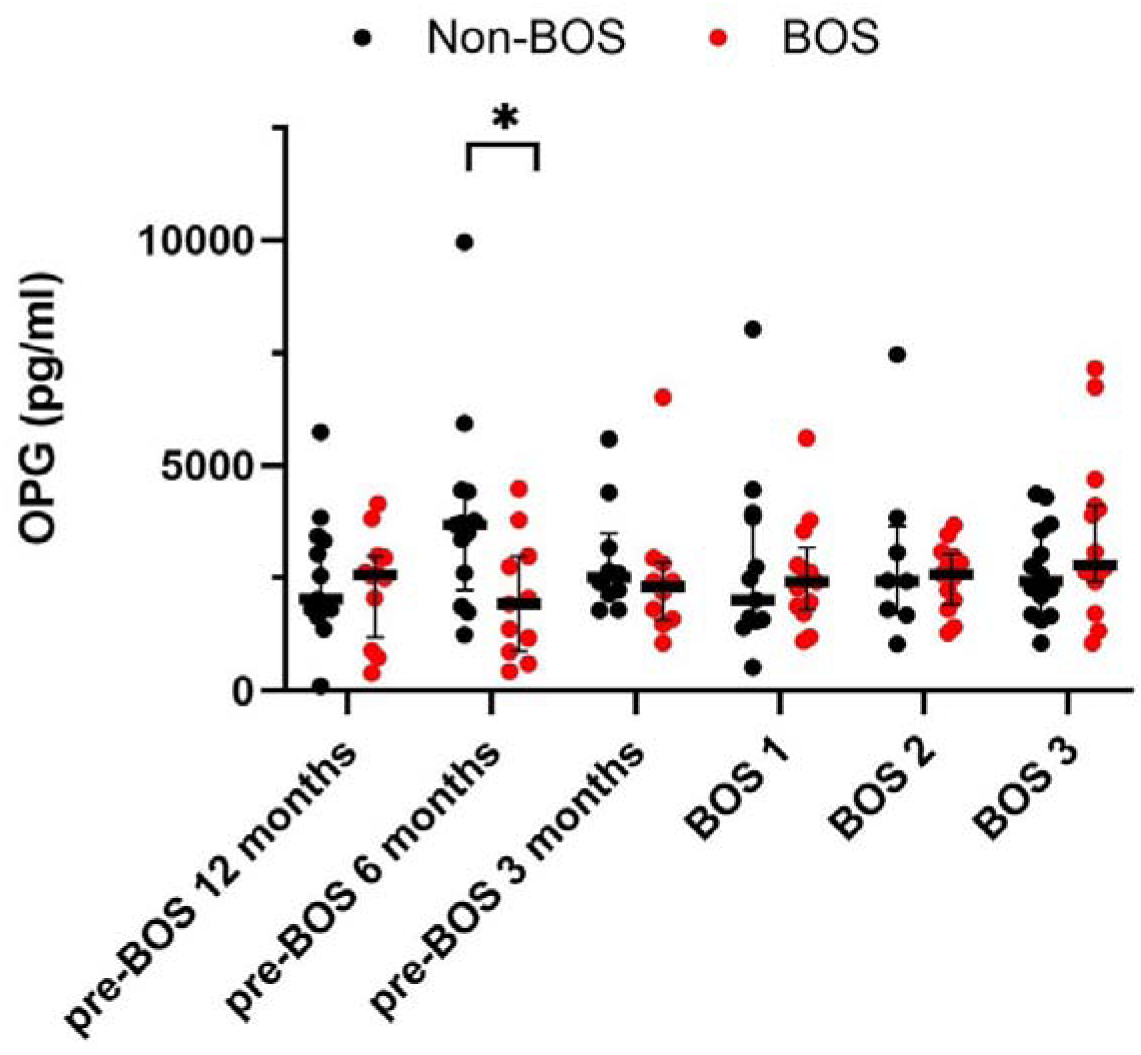
Osteoprotegerin (OPG) serum levels measured with ELISA in BOS and non-BOS patients (BOS n=19, non-BOS n=19). Mann-Whitney U was used to compare serum levels between groups at different timepoints. All error bars reflect median [interquartile range]. Longitudinal differences within BOS and non-BOS groups were analyzed with Wilcoxon-signed rank test. Abbreviations: BOS; bronchiolitis obliterans syndrome: BOS1; BOS stage 1: BOS2; BOS stage 2: BOS3; BOS stage 3. *p-value <0.05.

**Figure 6.**
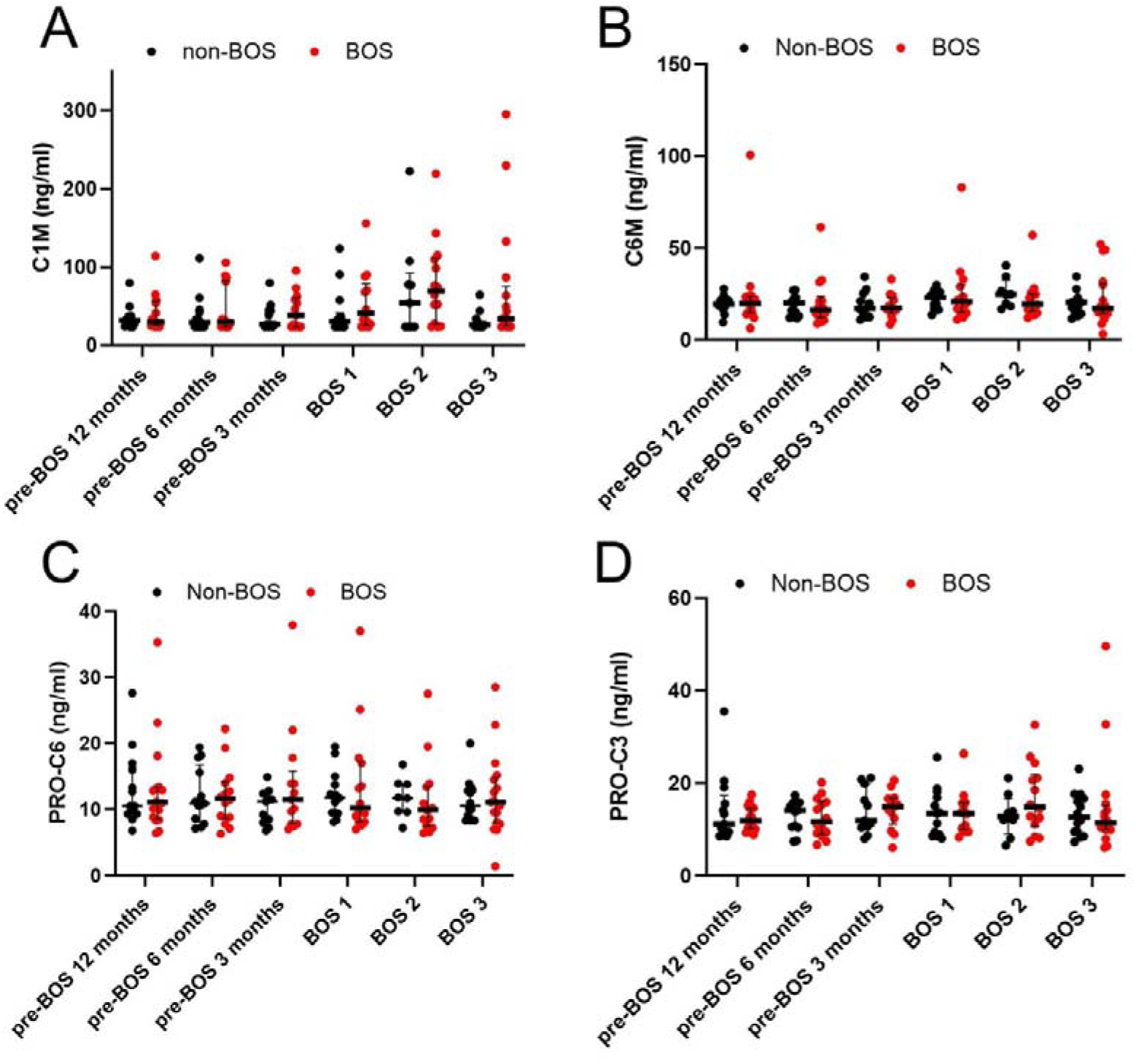
Serum levels of collagen fragments C1M, C6M, PRO-C3 and PRO-C6 at different timepoints for BOS and non-BOS patients (BOS n=19, non-BOS n=19). Differences between BOS and non-BOS patients were analyzed with Mann-Whitney U test at each timepoint. Longitudinal differences within BOS and non-BOS groups were analyzed with Wilcoxon-signed rank test. All error bars reflect median with interquartile range. Abbreviations: BOS; bronchiolitis obliterans syndrome: BOS1; BOS stage 1: BOS2; BOS stage 2: BOS3; BOS stage 3. C1M: type I collagen degradation; C6M: type VI collagen degradation; PRO-C3: type III collagen formation; PRO-C6: type VI collagen formation.

### 3.1.3 Proteomics analysis

To identify potentially new fibrotic and inflammatory markers for development of BOS we used a hypothesis free approach for serum protein profiling. Through LC-MS/MS, serum samples from BOS and non-BOS patients were analyzed three months before BOS onset, at BOS stage 1 and BOS stage 3, after depletion of most abundant proteins such as albumin.

### 3.1.4 NCHL1, K2C6A and CENF expression was decreased, while FANCE expression was increased in BOS patients compared to non-BOS patients 3 months before BOS

Three months before BOS onset, neural cell adhesion molecule L1-like protein (NCHL1), human type II Keratin-6 (K2C6A) and centromere protein F (CENPF) were lower in BOS patients compared to non-BOS patients (FDR<0.1, Figure 7, Table 2). Fanconi anemia complementation group E protein (FANCE) expression was higher in BOS patients compared to non-BOS patients (FDR<0.1, Figure 7, Table 2). However, these differences did not persist over time for any proteins, indicating a lack of stability of the protein markers.

**Figure 7.**
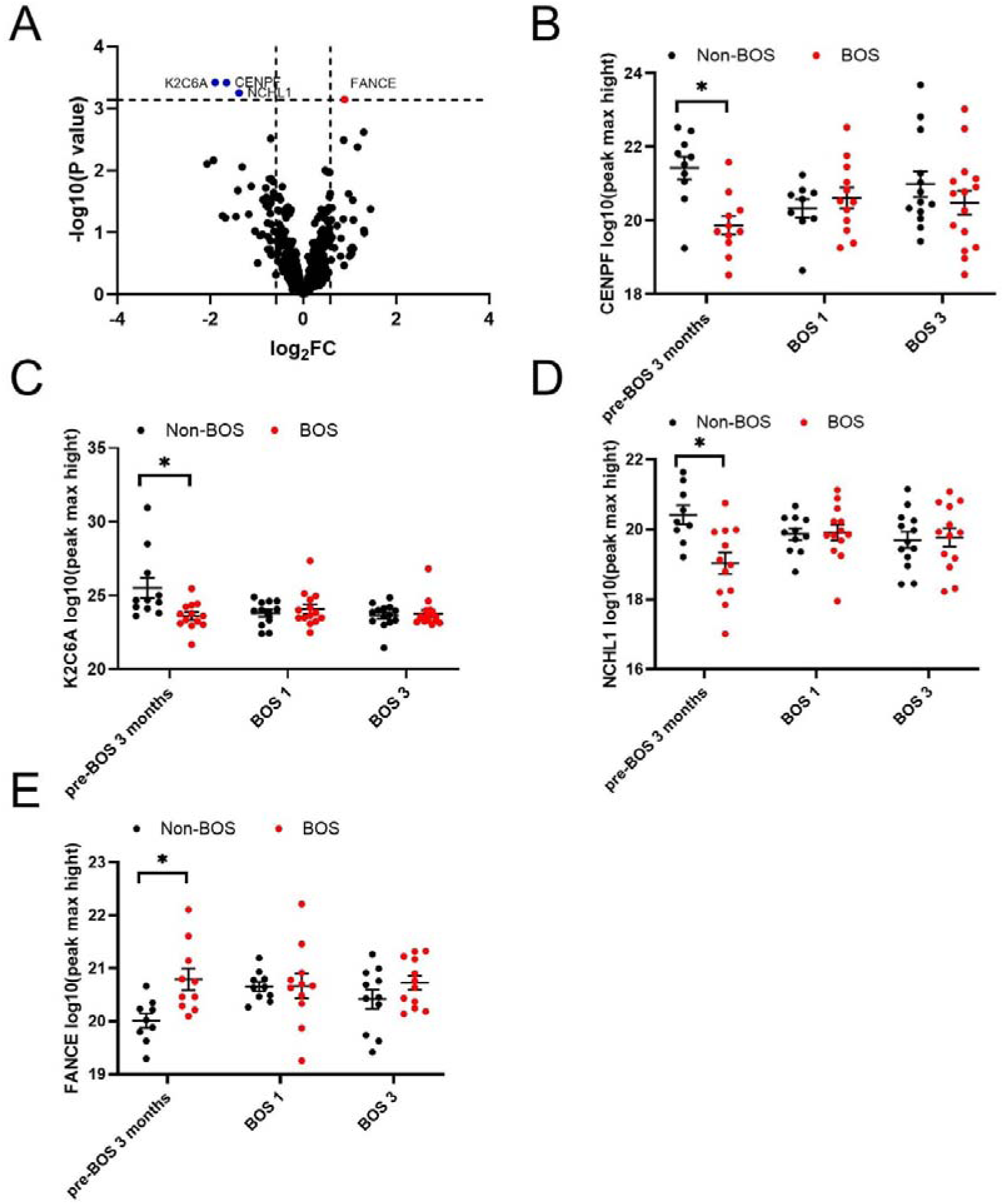
Proteins that were expressed at significantly different protein levels between BOS and non-BOS patients three months before onset of BOS (BOS n=18, non-BOS n=16). False discovery rate <0.1 was considered significant. Protein levels were log10 transformed. A: overview of significant differentially expressed proteins in proteomic analysis in BOS compared to non-BOS patients, red indicates overall higher expression, blue indicates lower expression in BOS compared to non-BOS patients. B: CENPF expression 3 months pre-BOS, at BOS stage 1 and BOS stage 3 in BOS compared to non-BOS patients. C: K2C6A expression pre-BOS, at BOS stage 1 and BOS stage 3 in BOS compared to non-BOS patients. D: NCHL1 expression pre-BOS, at BOS stage 1 and BOS stage 3 in BOS compared to non-BOS patients. E: FANCE expression 3 months pre-BOS, at BOS stage 1 and BOS stage 3 in BOS compared to non-BOS patients. All error bars reflect standard error of the mean (SEM). Abbreviations: BOS: bronchiolitis obliterans syndrome: pre-BOS; 3 months before BOS onset: BOS1; BOS stage 1: BOS3; BOS stage 3. CENPF: centromere protein F; K2C6A: human type II Keratin-6; NCHL1: neural cell adhesion molecule L1-like protein; FANCE: Fanconi anemia complementation group E protein. * p-value <0.05.

**Table 2.** Proteins differentially expressed at protein level data three months before onset of BOS compared to non-BOS patients. False discovery rate <0.1 was considered significant.

| Peptide | logFC | p-value | False discovery rate |
| --- | --- | --- | --- |
| **CENPF** | -1.65 | $3.83 \cdot 10^{-4}$ | 0.0813 |
| **K2C6A** | -1.90 | $3.78 \cdot 10^{-4}$ | 0.0813 |
| **NCHL1** | -1.38 | $5.62 \cdot 10^{-5}$ | 0.0813 |
| **FANCE** | 0.887 | $7.13 \cdot 10^{-4}$ | 0.0813 |

Abbreviations: logFC: log fold change; CENPF: centromere protein F; K2C6A: human type II Keratin-6; NCHL1: neural cell adhesion molecule L1-like protein; FANCE: Fanconi anemia complementation group E protein.

### 3.1.5 CRP expression is higher at BOS stage 3 in BOS compared to non-BOS patients

At BOS stage 3, CRP expression was higher in BOS patients compared to non-BOS patients at peptide level in LC-MS/MS analysis. Even though a trend towards higher expression of CRP was observed three months before BOS onset and at BOS diagnosis, this failed to reach significance in our limited dataset (Figure 8, panel A and B). For validation, CRP serum results from BOS and non-BOS patients were collected from electronic patient records to compare results to proteomics data. In serum, routinely collected CRP levels at end stage BOS were significantly higher in BOS patients compared to non-BOS patients (Figure 8, panel C, predicted difference 9.5 mg/L, 95% CI 1.6 - 17.4, p<0.01).

**Figure 8.**
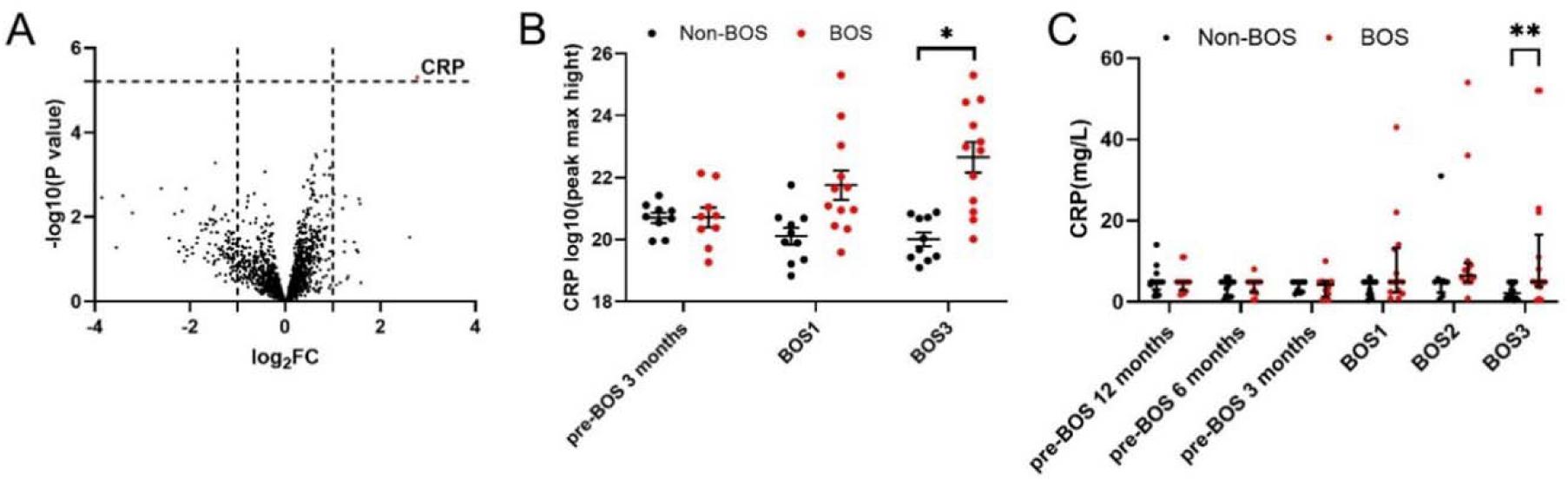
Protein that was expressed significantly different between BOS and non-BOS patients at peptide level data at BOS stage 3 in proteomic analysis and validated with retrospectively collected patient data. False discovery rate <0.05 was considered significant. A: An overview of significant differentially expressed protein in proteomic analysis in BOS compared to non-BOS patients, red indicates overall higher expression in BOS patients compared to non-BOS patients. B: Serum C-reactive levels measured with proteomic analysis in BOS compared to non-BOS patients 3 months before BOS onset, at BOS stage 1 and BOS stage 3. Peptide levels were log transformed to increase normality. C: For serum samples collected during regular follow up, CRP levels reported in mg/l in patient records were collected and analyzed with Mann-Whitney U (BOS n=17, non-BOS n=14). Error bars reflect standard error of the mean (SEM) in panel B, error bars reflect median with interquartile range in panel C. Abbreviations: CRP: C-reactive protein; BOS: bronchiolitis obliterans syndrome: pre-BOS; 3 months before BOS onset: BOS1; BOS stage 1: BOS3; BOS stage 3. * p<0.05 ** p< 0.01.

### 3.1.6 SERPINA3, CRP, PZP, ITIH3 and C4BP expression is higher at BOS stage 3 in BOS compared to non-BOS patients

Feature level analysis did not show significant differences for proteins in BOS patients compared to non-BOS patients before BOS onset. However, at BOS stage 3 several features were increased in BOS patients. These (Figure 9, Table 3) showed that SERPIN Family A member 3 (SERPINA3) and CRP had significantly higher expression in BOS patients compared to non-BOS patients (FDR<0.05). Also, Pregnancy Zone Protein (PZP), Inter-Alpha-Trypsin Inhibitor Heavy Chain 3 (ITIH3) and C4 binding protein (C4BP) expression were significantly higher at end stage BOS in BOS patients when compared to non-BOS patients (FDR<0.1).

**Figure 9.**
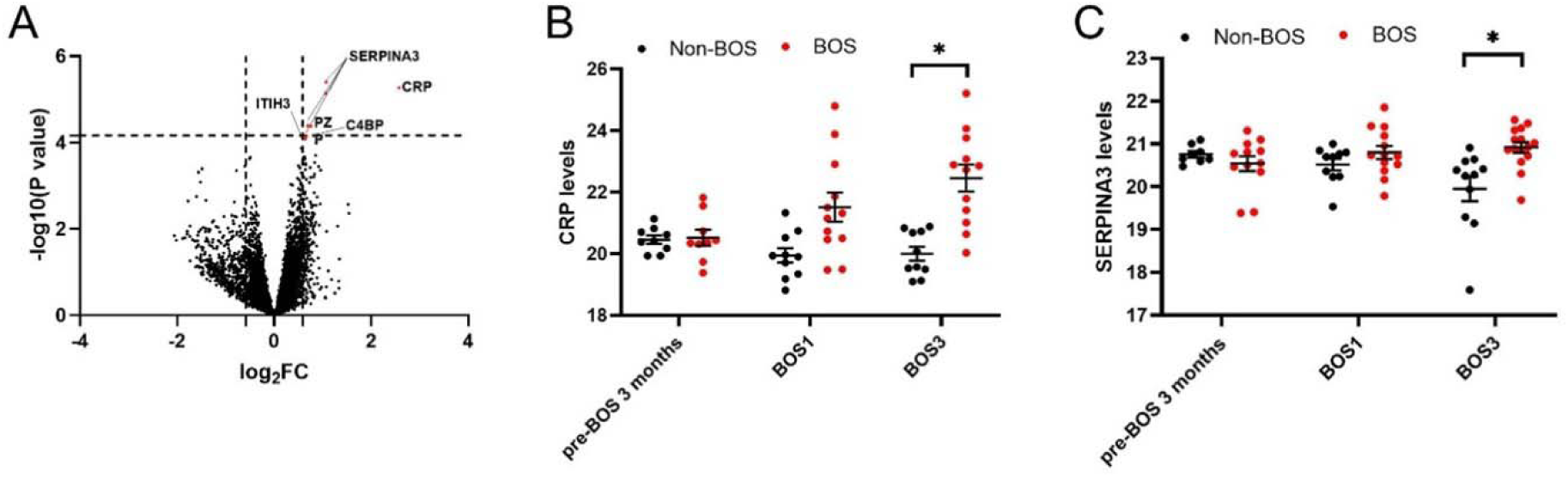
Significant proteins and their expression levels in BOS patients and non-BOS patients (BOS n=18, non-BOS n=16) at BOS stage 3 at feature level, analyzed from serum proteomics. False discovery rate <0.1 was considered significant. Peptide levels were log transformed to increase normality. Panel A: overview of significant differentially expressed peptides in proteomic analysis in BOS compared to non-BOS patients three months before BOS onset. Red indicates overall higher expression in BOS patients compared to non-BOS patients. Peptide levels were log transformed to improve normality. Panel B: Serum C-reactive peptide levels measured with proteomic analysis in BOS compared to non-BOS patients 3 months before BOS onset, at BOS stage 1 and BOS stage 3. C: SERPINA3 levels measured with proteomic analysis in BOS compared to non-BOS patients 3 months before BOS onset, at BOS stage 1 and BOS stage 3. All error bars reflect standard error of the mean (SEM). Abbreviations: CRP: C-reactive protein; SERPINA3: SERPIN A Family member 3 BOS: bronchiolitis obliterans syndrome: pre-BOS; 3 months before BOS onset: BOS1; BOS stage 1: BOS3; BOS stage 3. * p<0.05.

**Table 3.**
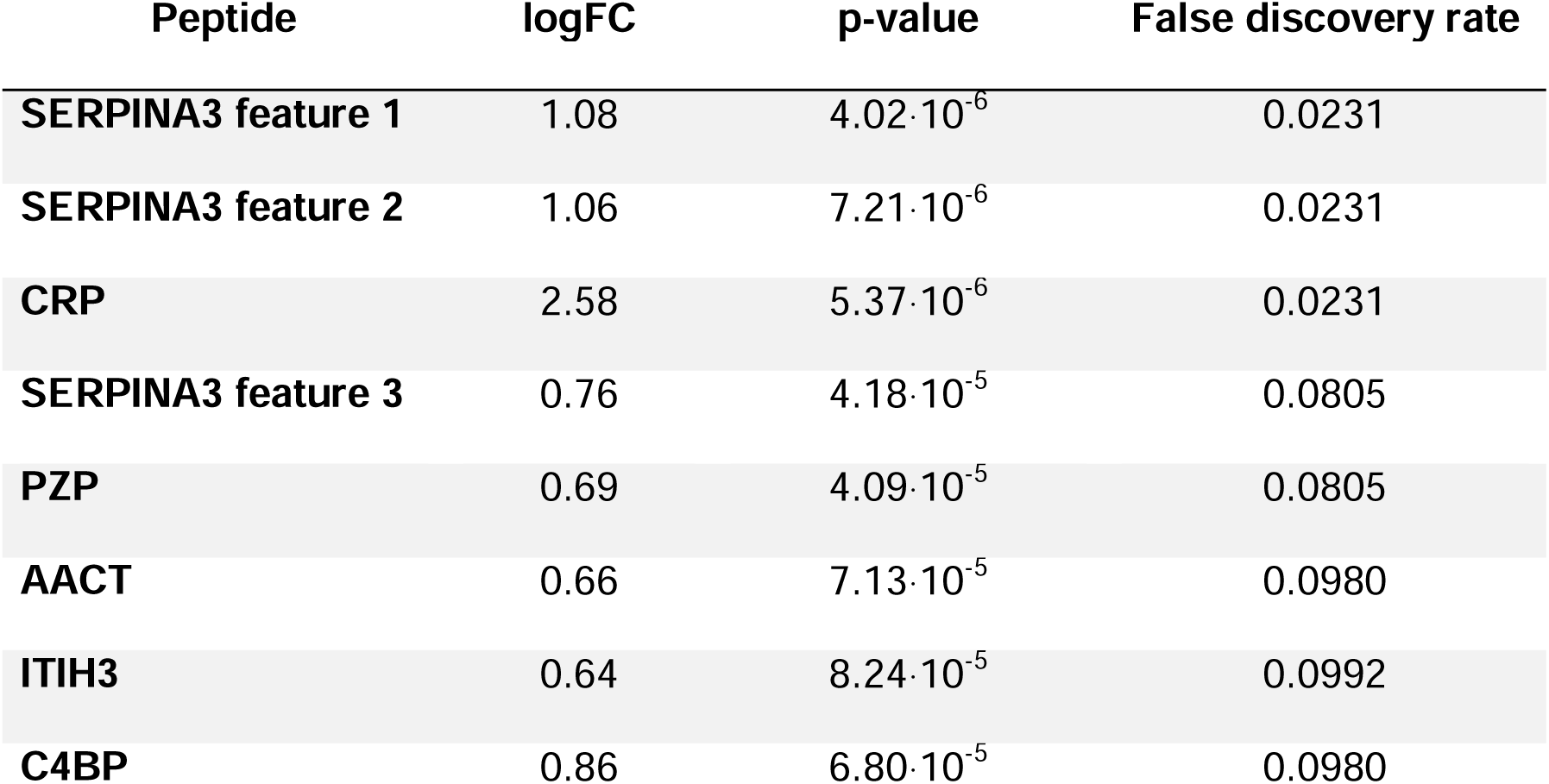
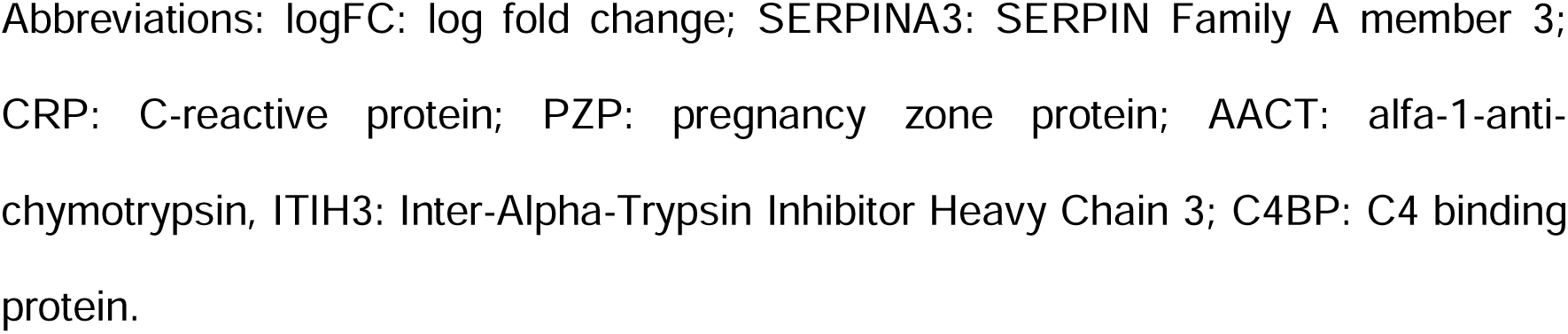
Protein expression at feature level with significant differential expression at BOS stage 3 in BOS patients compared to non-BOS patients. False discovery rate or adjusted p-value <0.1 was considered significant.

| Peptide | logFC | p-value | False discovery rate |
| --- | --- | --- | --- |
| SERPINA3 feature 1 | 1.08 | $4.02 \cdot 10^{-6}$ | 0.0231 |
| SERPINA3 feature 2 | 1.06 | $7.21 \cdot 10^{-6}$ | 0.0231 |
| CRP | 2.58 | $5.37 \cdot 10^{-6}$ | 0.0231 |
| SERPINA3 feature 3 | 0.76 | $4.18 \cdot 10^{-5}$ | 0.0805 |
| PZP | 0.69 | $4.09 \cdot 10^{-5}$ | 0.0805 |
| AACT | 0.66 | $7.13 \cdot 10^{-5}$ | 0.0980 |
| ITIH3 | 0.64 | $8.24 \cdot 10^{-5}$ | 0.0992 |
| C4BP | 0.86 | $6.80 \cdot 10^{-5}$ | 0.0980 |

Abbreviations: logFC: log fold change; SERPINA3: SERPIN Family A member 3; CRP: C-reactive protein; PZP: pregnancy zone protein; AACT: alfa-1-anti-chymotrypsin, ITIH3: Inter-Alpha-Trypsin Inhibitor Heavy Chain 3; C4BP: C4 binding protein.

### 4.1 Discussion

In this retrospective cohort study, we found that BOS patients exhibit both an increase in FANCE expression and decrease in K2C6A expression, suggesting an aberrant wound healing response, as well as elevated CRP expression, reflecting a low-grade inflammatory response.

Unexpectedly, serum levels of collagen neoepitopes did not differ between BOS and non-BOS patients. We hypothesized that OPG and collagen neoepitopes would be higher in BOS compared to non-BOS patients, as is seen in IPF^17,25,26^, but could not confirm this hypothesis in our small cohort. Possibly, OPG and collagen fragment levels in serum do not reflect the process of intraluminal fibrosis due to the inability of these markers to be released systemically from localized lesions, the hallmark of BOS. In comparison, the process of alveolar fibrosis in IPF patients is more diffuse and located in closer proximity to the circulation.. Lung tissue analysis with specific staining for these markers in BOS and non-BOS patients may shed a different light on the involvement of remodeling of these specific collagens.

NCHL1, K2C6A and CENPF protein expression were decreased three months before BOS onset, while FANCE expression increased. All failed to show persistence during the progression of BOS. Nevertheless, these findings shed new light on potential pathways involved in the early pathogenesis of BOS. NCHL1 is known to have tumor suppressive effects and is associated with prolonged survival in lung cancer ^27^. The role of NCHL1 in healthy lung is unknown. K2C6A also has a yet undefined role in lung cancer cell proliferation and migration^28,29^. Downregulation of CENPF, a mitotic cell regulator, could result in decreased cell proliferation^30^. FANCE is one of the eight proteins that form the core complex needed for monoubiquitination of the FANCD2-FANCI complex. This complex is needed for the intricate process of repairing DNA. Mutations or isoforms of one of the proteins of this core complex lead to increased risk of cancer in certain genetic diseases^31^. It is not known what an increase in FANCE expression in BOS reflects. In this study we did not analyze if increased FANCE expression in this cohort is for example due to a presence of an alternative isoform. The role of ITIH3, PZP and C4BP in the process of BOS and in end stage BOS remains to be elucidated. While ITIH3 is involved in stabilization of the extracellular matrix^32^, PZP and C4BP both serve as immunosuppressive proteins on for example T helper cells and the complement system^33–36^. To date, these proteins have not been described in (fibrotic) lung diseases.

We found higher expression of CRP and SERPINA3 in end stage BOS compared to non-BOS patients. CRP has multiple functions as an immunologic moderator. CRP stimulates phagocytotic cells, activation of the complement system, opsonization, and production of anti-inflammatory cytokines^37^, while also moderating the migration of fibroblasts^38^. CRP can increase up to 1000–fold in acute infection, however, low serum levels are related to low grade inflammation in rheumatological and cardiovascular disease^39–41^. Interestingly, the levels of CRP in BOS patients in our cohort remain relatively low and could be attributed to low grade inflammation. Vos et al showed that increased CRP plasma and bronchoalveolar lavage (BAL) fluid levels 90 days post-LTx were predictive of graft failure three years after LTx, underlining its clinical relevance^42,43^. SERPINA3 is a protease inhibitor, that prevents damage from neutrophil elastase released during inflammation^44^. Recently, SERPINA3 was identified to be higher in severe asthma patients compared to mild asthma patients and healthy controls^45^. Epithelial to mesenchymal transition, also a well-known pathogenic phenomenon in BOS, was enhanced by SERPINA3 in triple negative breast cancer and glioblastoma^46,47^. Future analysis of SERPINA3 lung tissue expression, BAL fluid levels collected prospectively and validation in a larger cohort in BOS and non-BOS patients’ serum would aid in further elucidating the role of SERPINA3 in BOS.

Even though differences in any of our identified proteins’ expression did not persist with progression of BOS, our results taken together might point towards aberrant repair, fibrogenesis and modifications in cell life cycles that precede BOS and increased inflammatory status at end stage BOS. The differentially expressed proteins such as CRP, FANCE and CENPF we identified in this study might refer to this process of increased inflammation, decreased resilience of cells, aberrant cell repair and increased apoptosis. We also hypothesize that within the complex pathogenesis of BOS, SERPINA3 could be a missing link in the interlude between the intertwined processes of inflammation and fibrosis in BOS due to its known role in epithelial mesenchymal transition and inflammation. Different cohorts including more patients with more timepoints before and after BOS onset to investigate the further significance of these proteins in BOS are needed. This study was the first study to perform label-free serum proteomics on several time points in BOS, yielding new possible markers and pathways to investigate. The historical availability of these sequential serum samples enabled us to perform these analyses on samples before and after onset of BOS, increasing sensitivity. By strictly matching BOS and non-BOS patients we aimed to decrease bias with respect to sex, age, use of immunosuppression and disease before LTx. However, this study does have some limitations. The main limitation of this study is that the current results do not establish a causal link between the markers found in the serum and the histological features characterizing BOS. Additional research, focusing on these markers in lung tissue is needed to further elucidate the role of these markers in BOS. The total number of patients included in this study is small due to the strict matching of BOS and non-BOS patients which does decrease the power of this study. Also, not all patients had serum samples available for all time points, because of the retrospective nature of this study, which could influence results. Furthermore, it is not known if other diseases of patients have influenced the results in this study, since measurements in serum are not restricted to diseases in the lung and patients were not matched for comorbidities. Ideally, we would have used BAL fluid to validate our results. Hypothetically, BAL fluid can reflect pulmonary tissue concentrations of serum markers more accurately and is less affected by systemic processes, However we had no availability to BAL fluid from these patients. Therefore, future validation of results is necessary in different media like BALf or lung tissue. Also, the staging for BOS severity changed in 2019, now also including stage 4 (FEV1<35% of baseline)^5^, however due to the historical nature of our data with inclusion until 2017, we decided to apply the earlier staging criteria of the ISHLT^18^.

## 5.1 Conclusion

In conclusion, this study suggests that several pathways are involved in the pathology of BOS, reflecting a complex interplay between both fibrosis and inflammation in BOS. The results should be further investigated in larger, prospective studies to expand the knowledge on their role in the development of BOS.

## Conflict of interest

EA van der Ploeg: none. A. Faiz: none. G. Teitsma: none. A. Sanchez Brotons: none, P. Horvatovich: none, N. Govorukhina: none. B.N. Melgert: none. J.M.B Sand and D.J. Leeming are employees of Nordic Bioscience and own stocks. J.K. Burgess: none, C.T.Gan: reports grants from Chiesi, via UMCG and from 2020 to 2024 participated in speaking activities, advisory committees and consultancies for Chiesi.

## Author contributions

Concept and design of this study were performed by EA van der Ploeg, A. Faiz, JK Burgess, B.N. Melgert, DJ Leeming, JMB Sand and CT Gan. Acquisition of the data was performed by G Teitsma, A Faiz, A. Sanchez Brotons, N. Govorukhina, DJ Leeming, JMB Sand and P. Horvatovich. Interpretation of the data was performed by EA van der Ploeg, A. Faiz, CT Gan and JK Burgess. Drafting of the manuscript was done by EA van der Ploeg and A.Faiz. G. Teitsma, P. Horvatovich, JK Burgess, A. Sanchez Brotons, N. Govorukhina, JMB Sand, DJ Leeming, BN Melgert and CT Gan critically revised the manuscript.

## Funding

This research was supported by a grant from Stichting Astma Bestrijding, grant number 2019/011 to C.T. Gan and Nederlandse Organisatie voor Wetenschappelijk Onderzoek (NOW) Aspasia premie subsidienummer 015.013.010 to J.K. Burgess.

## Data availability statement

All data generated or analyzed during this study are included in this article and its supplementary material files. Further enquiries can be directed to the corresponding author.

## Abbreviations

AATD: alpha-1 antitrypsin deficiency
BOS: bronchiolitis obliterans syndrome
C4BP: C4 binding protein
CLAD: chronic lung allograft dysfunction
CRP: C-reactive protein
FANCE: Fanconi anemia complementation group
E protein FEV1: Forced expiratory volume in one second
FVC: Forced vital capacity
ITIH3: Inter-Alpha-Trypsin Inhibitor Heavy Chain 3
K2C6A: human type II Keratin-6
LTx: lung transplantation
LC-MS/MS: liquid Chromatography with tandem mass spectrometry
MMF: mycophenolic acid
NCHL1: neural cell adhesion molecule L1-like protein
OPG: osteoprotegerin
PZP: Pregnancy Zone Protein
SERPINA3: SERPIN A family member 3

