## Supplementary files for "It takes two: Aberrant repair and low-grade inflammation characterizes bronchiolitis obliterans syndrome after lung transplantation in serum proteomic analysis"

*Table of contents*

Supplementary method	P 3

Supplementary figure 1	P 5

Supplementary table 1	P 7

Supplementary table 2	P 8

Supplementary table 3	P 9

*Supplementary method:*

*Proteomics analysis:*

Samples were depleted using PierceTM Top 2 Abundant Protein Depletion Spin Columns [Thermo Scientific]. The filtrate was collected by mixing the samples with the resin, using an end-over-end mixer for 30 minutes and then centrifuging the columns at 1000g for 2 minutes. The total protein concentration was determined using the PierceTM BCA Protein Assay Kit [Thermo Scientific].

Using liquid Chromatography with tandem mass spectrometry **(**LC-MS/MS) analysis, samples were injected on an ultrahigh performance nano liquid chromatography system (Dionex UltiMate 3000 RSLCnano, Thermo Scientific, Bremen, Germany) coupled to an Orbitrap mass spectrometer (Exploris 480, Thermo Fisher Scientific) with a nanoelectrospray source. The samples were loaded (2 µL/min) with the buffer A (0.1% formic acid (FA) in HPLC grade H2O) on a trapping column (Acclaim PepMap µ-precolumn, C18, 300 µm × 5 mm, 5 µm, 100 Ǻ, Thermo Scientific, Bremen, Germany). After sample loading, the trapping column was washed with 30 μL buffer A (3 μL/min), and the peptides were eluted (300 µL/min) onto a separation column (Acclaim PepMap 100, C18, 75 μm × 500 mm, 2 µm, 100 Ǻ, Thermo Scientific, Bremen, Germany). The spray was generated from a stainless-steel emitter (Proxeon, Thermo Scientifiec) at a capillary voltage of 1800 V. MS analyses were performed in positive ion mode and data-dependent acquisition mode (DDA). LC-MS/MS analysis was carried out in a cycle time of 1 s, and the dynamic exclusion duration was set to 30 s. An intensity threshold of 2·104, an isolation width of 0.7 m/z and an HCD collision energy of 30 NCE was used for MS/MS experiments. Precursor MS scans were performed over a m/z range from 350-1200, with a resolution of 120,000 FWHM at m/z 200 (RF Lens = 60%, maximum injection time= 247 ms, AGC target= 3·106) and MS/MS spectra were recorded with a resolution of 15,000 FWHM at m/z 200 with CID fragmentation and m/z range 100-1500 (maximum injection time= 40 ms, AGC target= 1·105).

All LC-MS/MS data were exported in mzML format using msconvert (version 3.0.11252) tool from ProteoWizzard toolset^47^. Peptide and protein identification was performed with philosopher^48^ and MSFragger^49–51^ using FDR 1% at peptide spectrum-matching, peptide and protein levels. For database search, human proteome from Swissprot was used. MS1 quantitative processing was performed with PASTAQ. For de novo protein identification, PEAKS was used.

*Supplementary figure*

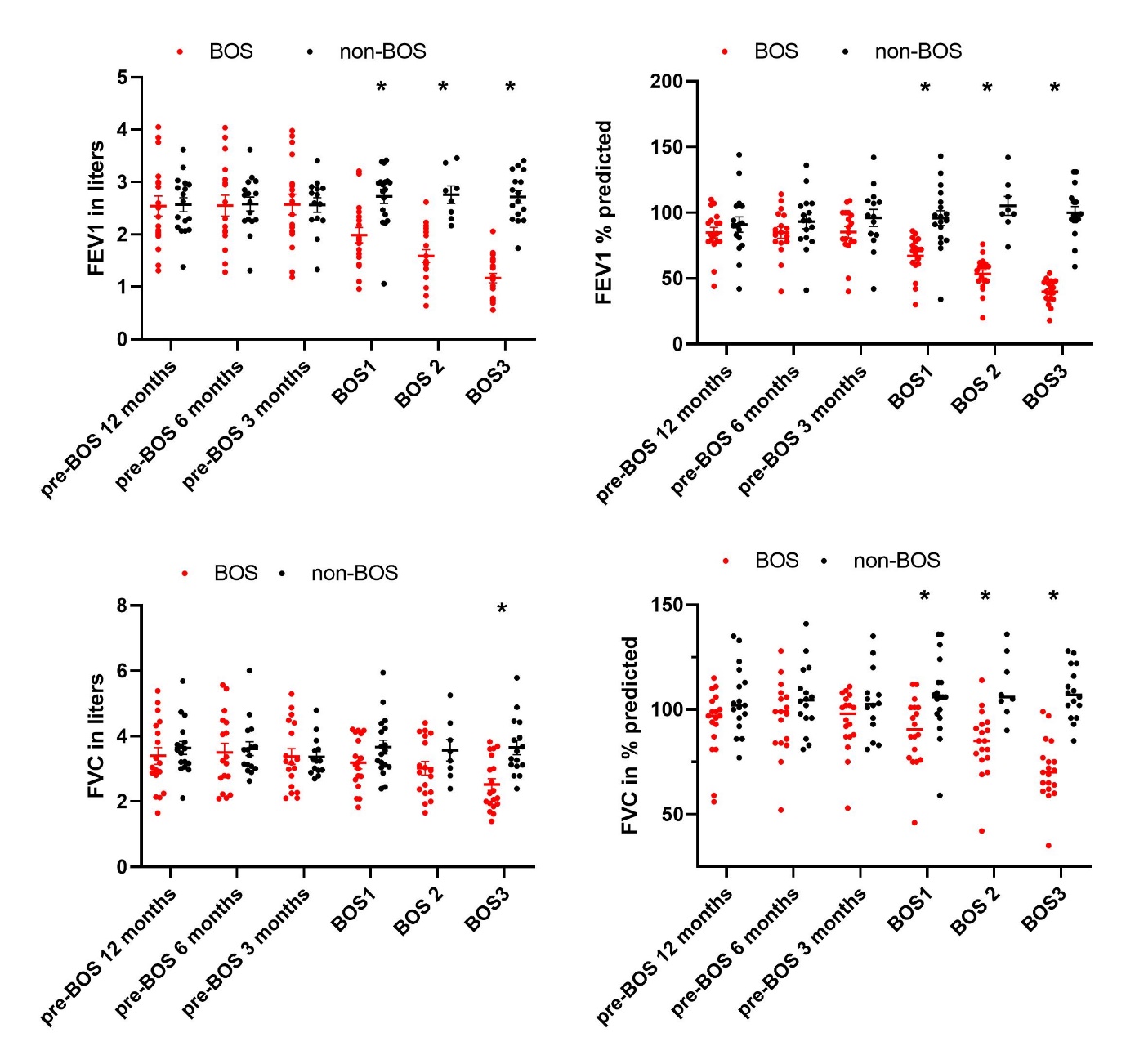

**Figure S1. Pulmonary function over time in BOS and non-BOS patients**. A: FEV1 in liters 12 months before BOS, 6 months before BOS, 3 months before BOS, at BOS stage 1, BOS stage 2 and stage 3 in BOS compared to non-BOS patients. B: FEV1 as a percentage of predicted 12 months before BOS, 6 months before BOS, 3 months before BOS, at BOS stage 1, BOS stage 2 and stage 3 in BOS compared to non-BOS patients. C: FVC in liters 12 months before BOS, 6 months before BOS, 3 months before BOS, at BOS stage 1, BOS stage 2 and stage 3 in BOS compared to non-BOS patients. D: FVC as a percentage of predicted 12 months before BOS, 6 months before BOS, 3 months before BOS, at BOS stage 1, BOS stage 2 and stage 3 in BOS compared to non-BOS patients. *p<0.05. All error bars reflect standard error of the mean (SEM). Abbreviations: BOS; bronchiolitis obliterans syndrome: BOS1; BOS stage 1: BOS2; BOS stage 2: BOS3; BOS stage 3

*Supplementary tables:*

**Table S1. Number of serum samples of BOS and non-BOS patients available for ELISA analysis for osteoprotegerin and collagen fragments.**

| **Number of serum samples included per ELISA analysis** | | | | | | | | | | | | |
| --- | --- | --- | --- | --- | --- | --- | --- | --- | --- | --- | --- | --- |
|  | -12 months before BOS  (n) | | -6 months before BOS  (n) | | -3 months before BOS  (n) | | BOS stage 1  (n) | | BOS stage 2  (n) | | BOS stage 3  (n) | |
|  | BOS | Non-BOS | BOS | Non-BOS | BOS | Non-BOS | BOS | Non-BOS | BOS | Non-BOS | BOS | Non-BOS |
| OPG | 12 | 12 | 11 | 13 | 10 | 10 | 13 | 13 | 13 | 8 | 15 | 14 |
| Collagen fragment |  |  |  |  |  |  |  |  |  |  |  |  |
| C1M | 16 | 13 | 14 | 13 | 13 | 11 | 13 | 14 | 14 | 9 | 17 | 13 |
| C6M | 16 | 13 | 14 | 13 | 13 | 11 | 14 | 13 | 14 | 8 | 17 | 14 |
| PRO-C6 | 16 | 14 | 14 | 14 | 13 | 11 | 14 | 14 | 14 | 8 | 17 | 14 |
| PRO-C3 | 16 | 13 | 14 | 13 | 13 | 11 | 14 | 14 | 14 | 8 | 16 | 14 |

Abbreviations: n: number; BOS: patients with bronchiolitis obliterans syndrome after lung transplantation; non-BOS: patients without bronchiolitis obliterans syndrome after lung transplantation; OPG: osteoprotegerin, BOS 1: BOS stage 1 (FEV1 66-80% of baseline), BOS 2: BOS stage 2 (FEV1 50-66% of baseline), BOS 3: BOS stage 3 (FEV1 <50% of baseline).

**Table S2. Number of serum samples of BOS and non-BOS patients available for proteomics analysis.**

| Patient group | -3 months before  BOS onset (n) | BOS stage 1 (n) | BOS stage 3 (n) |
| --- | --- | --- | --- |
| BOS (n=18) | 13 | 14 | 16 |
| Non-BOS (n=16) | 11 | 12 | 14 |

Abbreviations: n: number; BOS: patients with bronchiolitis obliterans syndrome after lung transplantation; non-BOS: patients without bronchiolitis obliterans syndrome after lung transplantation; BOS 1: BOS stage 1 (FEV1 66-80% of baseline), BOS 2: BOS stage 2 (FEV1 50-66% of baseline), BOS 3: BOS stage 3 (FEV1 <50% of baseline).

**Table S3. Time in days between lung transplantation and collection of serum sample (median [IQR]) in BOS and non-BOS patients according to timepoint**. Mann-Whitney-U test was used for analysis between groups.

| **Timepoint serum sample** | **BOS** | **Non-BOS** | **p-value** |
| --- | --- | --- | --- |
| -12 months before BOS onset - days | 650 [351-1749] | 739 [395 -1754] | 0.9 |
| -6 months before BOS onset - days | 977 [486-1951] | 857 [570-2014] | 1.0 |
| -3 months before BOS onset - days | 950 [701-2140] | 928 [622-1689] | 0.7 |
| BOS stage 1 - days | 1032 [839 -2121] | 1023 [651-1790] | 1.0 |
| BOS stage 2 - days | 1087 [723-1828] | 1100 [767-1898] | 1.0 |
| BOS stage 3 - days | 1812 [906-2546] | 1157 [860-2525] | 1.0 |

Abbreviations: BOS: patients with bronchiolitis obliterans syndrome after lung transplantation; non-BOS: patients without bronchiolitis obliterans syndrome after lung transplantation.
